# Sir proteins impede, but do not prevent, access to silent chromatin in living *Saccharomyces cerevisiae*

**DOI:** 10.64898/2025.12.02.691921

**Authors:** Kenneth Y. Wu, Zhuwei Xu, Hemant K. Prajapati, Peter R. Eriksson, David J. Clark

## Abstract

Gene silencing at the mating type loci in budding yeast (*HMRa* and *HMLα*) depends on the Sir proteins. Sir2, Sir3 and Sir4 are indispensable for silencing, whereas Sir1 has a more limited role. The Sir proteins are also involved in repression at telomeres and ribosomal DNA (rDNA) repeats. Proposed mechanisms for Sir-mediated silencing include limiting access to silenced DNA and inhibition of transcript initiation and/or elongation. Using an inducible M.SssI DNA methyltransferase expression system, we showed previously that the silenced mating type loci are methylated at a much slower rate than the rest of the genome in vivo. Here, we show that, in the absence of Sir2, Sir3 or Sir4, the silenced loci and the telomeric X elements are methylated at a similar rate to the rest of the genome, indicating that these Sir proteins impede access, but do not prevent it. The rDNA repeats are methylated faster in the absence of Sir2 and, to a lesser extent, of Sir3, but not of Sir4. The methylation rates of adjacent rDNA repeats are not strongly correlated in wild-type cells, suggesting that Sir proteins silence rDNA repeats randomly. Loss of Sir1 increases the methylation rate at *HMRa* but not at *HMLα*, telomeres or rDNA. Our data indicate that steric occlusion is unlikely to be the primary mechanism of silencing, since silenced DNA is accessible in vivo, albeit at a slower rate than elsewhere in the genome.

---

DNA in eukaryotes associates with histones and non-histone proteins to form chromatin. Chromatin can adopt active states permissive to transcriptional activity or silent states restrictive to transcriptional activity, the regulation of which is vital for proper growth and development in eukaryotic organisms. The process of establishment, maintenance and inheritance of such silent chromatin states is known as silencing (1).

In the budding yeast *Saccharomyces cerevisiae*, silent chromatin domains are found at the cryptic homothallic mating (*HM*) loci *HMLα* and *HMRa*, certain telomeric regions, and ribosomal DNA elements (rDNA) (reviewed by (2, 3)). At the *HM* loci, silencing is controlled by sequence-specific silencer elements, which flank the region and contain at least two of three possible binding sites for Abf1, Rap1 or Orc1 proteins. At the silencers, the role of these factors is to facilitate the binding of the <u>s</u>ilent information regulator (Sir) proteins, all of which are necessary for the stability of the silent state. Sir1 is recruited to the silencer by the silencer-bound proteins and in turn recruits Sir2, Sir3 and Sir4. Sir2 preferentially, but not exclusively, deacetylates lysine 16 of histone H4 (H4K16), Sir3 binds to nucleosomes with unacetylated H4K16 residues, and Sir4 directly binds the other Sir proteins and serves as the scaffolding. H3 and H4 in adjacent nucleosomes are deacetylated by Sir2, facilitating the binding and spreading of more Sir2/3/4 complexes along the locus, which is thought to sterically hinder transcriptional activator and general transcriptional machinery occupancy within the silent domain (4). Sufficiently strong activating regulatory elements can nevertheless overcome this transcriptional repression, and this effect can be in turn counteracted by recruitment of more Sir proteins (5). Altogether, these data suggest that the silencing machinery competes with the transcriptional machinery for DNA access.

Sir1 recruitment is not required for silencing at the telomeric regions. At the nuclear periphery, Orc1 and Abf1 recruit Sir2, Sir3 and Sir4 to the X elements (6), while Rap1 recruits Sir2, Sir3 and Sir4 to the telomere ends (7–9), where they are thought to contribute to nuclear organization and facilitate a protective structural folding of the telomere ends back to the X elements. One proposed function of X element silencing is that it inhibits the expression of telomeric repeat-containing RNAs, which form DNA-RNA hybrids (R loops) that stall the replication fork and cause DNA double-strand breaks that facilitate telomere elongation (10).

In addition to Sir1, Sir3 and Sir4 are also dispensable for silencing at the rDNA (11). The nucleolus contains 150 to 200 tandem 9.1 kb rDNA repeats, around half of which are transcriptionally active (12–14). Each repeat includes a 35S rRNA gene transcribed by RNA polymerase I (Pol I) that is later processed to form mature rRNAs, and a 5S rRNA gene transcribed by RNA polymerase III (Pol III) flanked by <u>n</u>on-<u>t</u>ranscribed <u>s</u>pacers, *NTS1* and *NTS2*. At *NTS1*, the RENT complex, which contains Sir2, Cdc14 (15) and Net1 (16), is critical for repressing E-pro, a promoter driving the expression of non-coding RNA polymerase (Pol II) transcripts. E-pro promotes the dissociation of cohesin responsible for inhibiting rDNA copy number changes driven by mitotic recombination (17).

Various molecular probes have been used to analyze silent chromatin accessibility in budding yeast. Generally, these probes have been used on the principle that nucleosomes prevent access to their DNA. Dam methyltransferase, which methylates adenine in exposed GATC sites, has been used to probe genomic loci in living yeast cells, including at the mating loci (18) and telomeres (19). Loo and Rine used both HO endonuclease, which catalyzes mating type switching of the *MAT* locus, and restriction enzymes, in early in-vitro assays for *HM* accessibility in nuclei (20). The Simpson group, on the other hand, used micrococcal nuclease (MNase) to map nucleosome positions at *HMLα* (21) and *HMRa* (22) in isolated nuclei. In a more recent study, Brothers and Rine used Sir-EcoGII fusion proteins to map association of the Sir proteins (23). Our own study used a genomic approach with both Dam and M.SssI methyltransferases to quantitatively assay DNA accessibility in living yeast cells. We demonstrated that the yeast genome is globally accessible, even in α-factor arrested cells, except for the centromeres and the *HM* domains (24, 25). Here, we extend our analysis to *sirΔ* mutants in vivo, shedding light on how silent domains achieve transcriptional repression in the context of a dynamic chromatin environment.

## Results

To investigate the contributions of the Sir proteins to the accessibility of the silent chromatin state in living yeast cells, we created *sir1Δ, sir2Δ*, *sir3Δ* and *sir4Δ* strains with a cassette encoding M.SssI, which is a bacterial 5mC methyltransferase targeting CG sites (Fig. 1A). *M.SssI* is driven by a promoter containing a high-affinity Gcn4 binding site (24, 26). When these cells are cultured in medium without isoleucine and valine, the addition of sulfometuron methyl (SM) induces the production of Gcn4 (27), which in turn induces the expression of *M.SssI*. In this study, we used M.SssI rather than the Dam methyltransferase we used previously (24), which methylates GATC sites, because there are too few GATC sites in the mating type loci. We note that Dam methylates the yeast genome almost completely during the time course, whereas M.SssI is not expressed at sufficiently high levels to complete methylation (24). We did not utilize the auxin-dependent degron fused to M.SssI (Fig. 1A) because the background methylation levels were acceptably low (Fig. 1B). Cells were grown to log phase and then induced with SM. Samples were taken immediately before SM induction and at 30, 60, 120 and 240 min after induction. Successful induction was confirmed by immunoblotting for the 3HA tag fused to M.SssI using tubulin as a loading control (Fig. S1). Expression levels of M.SssI are similar in wild-type and all four *sirΔ* strains at each time point.

**Fig. 1.**
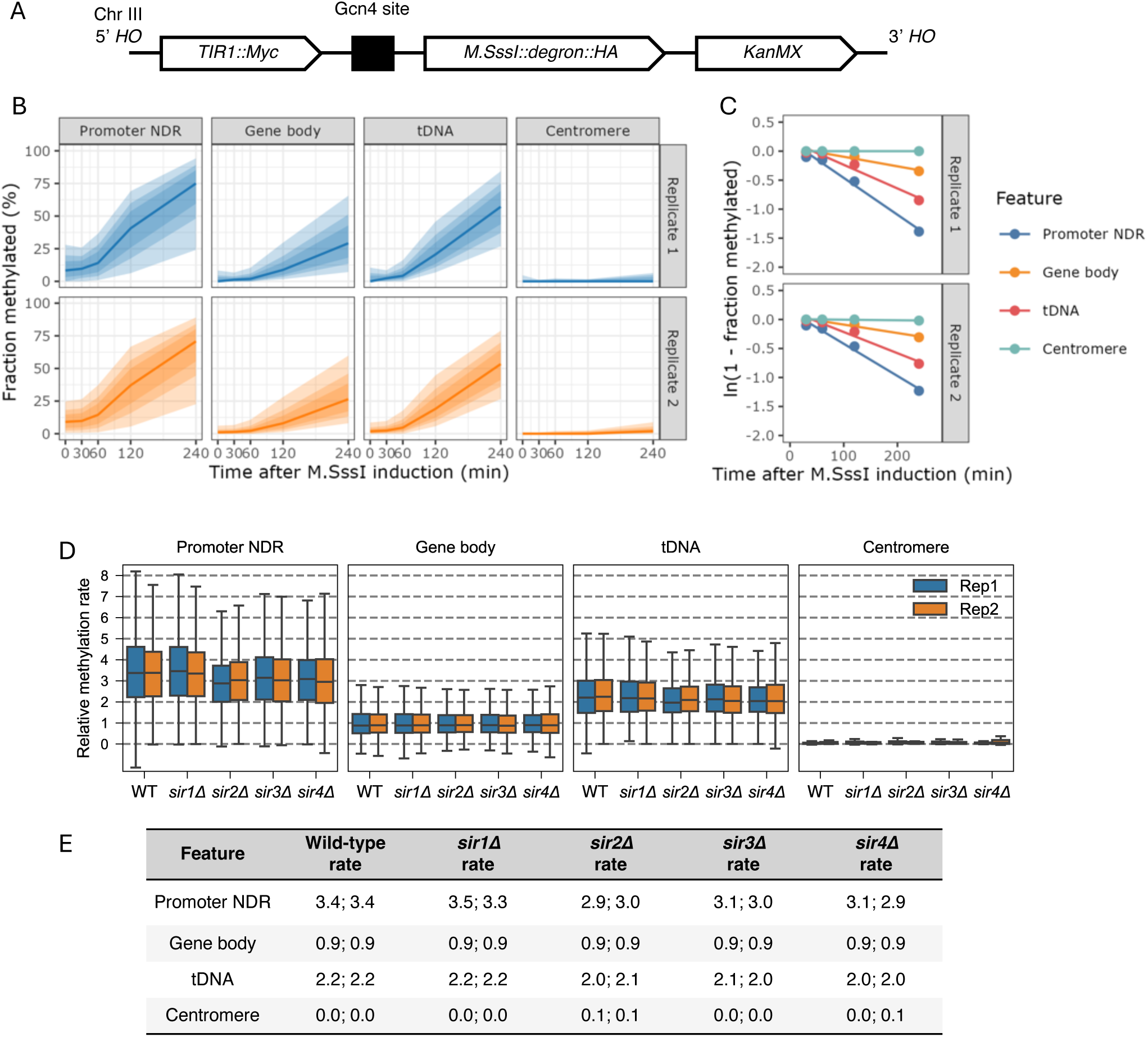
Genomic DNA in wild type, *sir1Δ*, *sir2Δ*, *sir3Δ* and *sir4Δ* strains is methylated at similar rates in vivo. (*A*) Schematic of the *M.SssI* construct. (*B*) Plots of the median methylated fraction of CG sites (solid line) for various genomic features against time after M.SssI induction. Shading: lightest to darkest: 5-95%, 15-85%, and 25-75% of all CG sites in the feature. (*C*) Genomic features are methylated at different rates. For each feature, the natural log of the median unmethylated fraction is plotted against time. The slope of the regression line is the methylation rate constant. (*D*) Box plots of methylation rate constants for each feature normalized to the whole-genome median (each box contains 25 to 75% of the data, the line is the median, and the whiskers represent 1.5 times the interquartile range to the farthest data points). (*E*) Median methylation rate constants for each feature normalized to the whole-genome median. Values from two biological replicate experiments are presented.

### Methylation rates of non-silenced chromatin regions

Aliquots of cells were collected at each time point, from which genomic DNA was purified and subjected to nanopore long read sequencing, which can distinguish 5mCG from CG (28). For each CG site in the genome, we calculated the fraction of reads in which the site is methylated. CG sites were grouped according to their location with respect to features of interest, and the median methylated fraction and interquartile ranges of these features were plotted over time. In wild-type cells, as we observed previously (24), promoter NDRs and tRNA genes are methylated faster than gene bodies, whereas centromeres are almost completely protected from methylation (Fig. 1B). Methylation rates were quantified by assuming pseudo-first order reaction kinetics (24). Rate constants for each region were derived from the slope of the regression line in plots of the natural log of the median unmethylated fraction against time (Fig. 1C).

The median relative rate and interquartile ranges of the data for all individual CG sites in each region are presented as box plots (Fig. 1D) with median rate constants (Fig. 1E). Rate constants were normalized to the genomic median rate constant (set at 1.0), which assumes that the genomic methylation rate is generally unaffected by loss of a Sir protein (i.e., that Sir proteins act at only a few genomic sites) and normalizes for small differences in M.SssI induction among the strains (Fig. S1). In wild-type cells, the median promoter NDR is methylated 3.4 times faster than the genomic median, whereas gene bodies are methylated slightly slower than the genomic median (relative rate = 0.9) and centromeres are almost completely protected (Fig. 1D,E). In the *sirΔ* strains, the rate constants for promoter NDRs, gene bodies and centromeres are very similar to wild type (Fig. 1D,E).

We confirmed that the chromatin organization on genes is generally unaffected in *sirΔ* mutants (Fig. S2). We generated methylation phasing profiles from the aggregate methylation rates at all ∼5,000 genes aligned on the dyad of the wild type +1 nucleosome for each of our strains. We compared them to the wild-type nucleosome dyad phasing profile from our previously obtained MNase-seq data (29). The latter plots represent normalized counts of the number of times each nucleotide is the central nucleotide in a nucleosomal DNA sequence protected from MNase digestion (the central nucleotide is equivalent to the nucleosome dyad). Accordingly, the two plots are out of phase, as methylation is faster in linkers and NDRs, whereas MNase-seq measures DNA regions protected from digestion. Note that MNase digestion is performed in nuclei, in which the chromatin is static and nucleosomes protect their DNA from both MNase and methylation, whereas in vivo, chromatin is dynamic, such that nucleosomal DNA can be methylated, albeit at a slightly slower rate than linkers (Fig. S2 (24)). The phasing profiles in the *sirΔ* mutants are very similar to that of the wild type.

### Methylation is slow at the silent *HM* loci in wild-type cells

In budding yeast, the genes located at the *MAT* locus determine the mating type of the cell, whereas the silent genes at *HML* and *HMR* serve as templates for the mating type switch mechanism that is absent in laboratory strains. As the strains used in this study are mating type *a* (*MATa*), the corresponding silent copy resides at *HMRa*, and the genetic information of the opposite mating type, *α,* resides at *HMLα*. Nanopore reads are sufficiently long to differentiate between the identical ∼1.6 kb sequences at *MATa* and *HMRa*.

In wild-type cells, both silenced loci are methylated more slowly than the active *MATa* locus (Fig. 2A, top panel). Each column represents the relative methylation rate at a specific CG site in the locus. There is some variation in methylation rate across each locus, particularly near the flanking E and I silencers at *HMLα* and *HMRa*, where the rates are fast. Since the median rate at *MATa* is 1.5 to 1.6 times the genomic median and the median rates at *HMLα* and *HMRa* are 0.4 to 0.5 times the genomic median (Fig. 2B,C), *MATa* is methylated ∼3 times faster than *HMLα* and *HMRa* in wild-type cells. Although previous studies have determined that *HMRa* silencer strength is higher than that of *HMLα*, irrespective of the enhancer and promoter driving transcription (30), this difference is not reflected in the median methylation rate constants between the two loci.

**Fig. 2.**
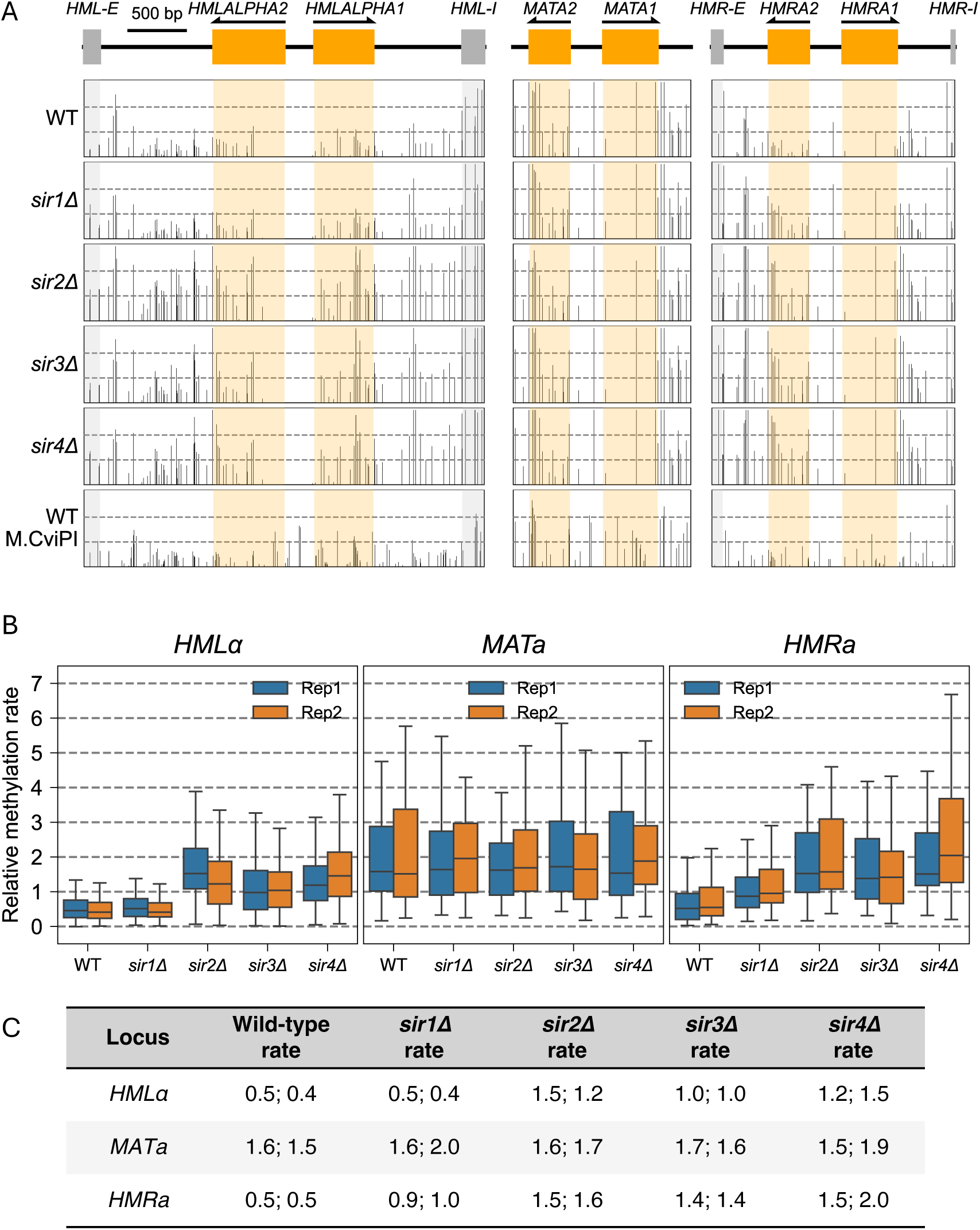
The silenced *HMLα* and *HMRa* loci are methylated faster in the absence of Sir2, Sir3 or Sir4 in vivo. (*A*) Methylation rate constants of CG sites in the *HMLα*, *MATa* and *HMRa* loci in wild-type, *sir1Δ*, *sir2Δ*, *sir3Δ* and *sir4Δ* strains. The lower dashed line represents the genomic median normalized to 1.0, while the upper dashed line represents twice the genomic median. (*B*) Box plots showing the distributions of the individual methylation rate constants of all CG sites in the *HMLα*, *MATa* and *HMRa* loci. Data for two biological replicate experiments are presented. Each box contains 25 to 75% of the data, the line is the median, and the whiskers represent 1.5 times the interquartile range to the farthest data points. (*C*) Median methylation rate constants for *HMLα*, *MATa* and *HMRa* normalized to the whole-genome median (set at 1.0). Data from two biological replicate experiments are presented.

CG sites are scarce in the divergent *HM/MAT* promoters: there is only one CG site in the *HMLα* promoter and only two CG sites in the *MATa/HMRa* promoter (Fig. 2A). To obtain data for these promoters and to confirm our observations with M.SssI using another DNA methyltransferase as a probe, we constructed a wild-type strain expressing M.CviPI instead of M.SssI. M.CviPI methylates C in GC sites (31, 32), and there are four or five GC sites in each promoter. The silenced promoter at *HMRa* is methylated more slowly than the identical active promoter at *MATa*, as expected (Fig. 2A, bottom panel). We used a longer time course because M.CviPI is induced more slowly than M.SssI (Fig. S3A). M.CviPI yields similar results to M.SssI: centromeres are essentially protected from methylation, gene bodies are methylated at a rate close to the genomic median, whereas promoter NDRs and tRNA genes are almost completely methylated (Fig. S3B). The relative rates are similar to those for M.SssI in wild-type cells (Fig. S3C,D). Importantly, we observed similar relative rates for *MATa* and the *HM* loci: *MATa* is methylated ∼1.3 times faster than the genomic median and ∼3 times faster than *HMRa* and *HMLα* (Fig. S3C,D). In conclusion, the silenced loci are methylated much more slowly than the active *MATa* locus in vivo by both M.SssI and M.CviPI.

### Sir2, Sir3 and Sir4 impede methylation at the silent *HM* loci

In the absence of Sir2, Sir3 or Sir4, *HMLα* and *HMRa* are methylated much faster by M.SssI at most CG sites within *HMLα* and *HMRa* (Fig. 2A). A useful direct comparison is between the *MATa* locus and the identical region in *HMRa* (drawn to the same scale): the CG sites in *HMRa* are methylated ∼3 times slower than the same sites in *MATa* in wild-type cells whereas, in *sir2Δ*, *sir3Δ* and *sir4Δ* cells, both loci are rapidly methylated (Fig. 2B). In more detail (Fig. 2C), the median relative methylation rate at *HMLα* is slow in wild-type cells (0.5; replicate: 0.4) and fast in *sir2Δ* cells (1.5 and 1.2). The rate at *HMRa* is also slow in wild-type cells (0.5 and 0.5) and fast in *sir2Δ* cells (1.5 and 1.6). The same is true of *sir4Δ* cells: median methylation rates at *HMLα* (1.2 and 1.5) and *HMRa* (1.5 and 2.0) are fast relative to wild-type cells (Fig. 2C). In *sir3Δ* cells, the median methylation rate at *HMRa* (1.4 in both replicates) is slightly slower than in *sir2Δ* and *sir4Δ* cells. Although methylation at *HMLα* (1.0 in both replicates) is slower in *sir3Δ* cells than in *sir2Δ* and *sir4Δ* cells, it is still about twice as fast as in wild-type cells (0.5 and 0.4). For comparison, the median rate at *MATa* is about the same in wild-type (1.6 and 1.5) and the *sir2Δ, sir3Δ* and *sir4Δ* mutants (ranging from 1.5 to 1.9) (Fig. 2C). We conclude that loss of Sir2, Sir3 or Sir4 results in increased methylation rates at the *HM* loci, implying that Sir2, Sir3 and Sir4 impede methylation in living cells.

### Sir1 impedes methylation at *HMRa* but not at *HMLα*

In the absence of Sir1, *HMLα* is methylated about as slowly as in wild-type cells, but *HMRa* is methylated at a rate intermediate between wild type and the other *sirΔ* mutants (Fig. 2B,C). This observation indicates that Sir1 impedes methylation at *HMRa* but has no effect at *HMLα*. It has been shown that *sir1Δ* cultures are a mixture of silenced and non-silenced cells (33–35). In particular, *HMRa* is de-repressed in a higher proportion of *sir1Δ* cells (∼90%) than *HMLα* (∼40%) (33, 35), which is consistent with our observation that *HMRa* is methylated faster than *HMLα* in *sir1Δ* cells. In summary, Sir2, Sir3 and Sir4 impede access to DNA at the silent mating loci, but Sir1 only affects access at *HMRa*.

### Sir2, Sir3 and Sir4, but not Sir1, impede methylation at telomeric X elements

In the Saccharomyces Genome Database (SGD), telomeres are defined as the region encompassing the ∼460 bp X element to the chromosome end, which varies from 829 bp to 35 kb in our YDC111 strain background. An X element is present on each arm of each chromosome. However, only a subset of regions within telomeres is subject to Sir-mediated silencing, including the X element and certain sub-telomeric genes (36). The telomeric regions, as defined in SGD, are methylated at a median rate close to the genomic median in wild-type cells (24) and in all the *sirΔ* mutants (Fig. 3A,B). Unlike telomeres as a whole, X elements are methylated at only about half the genomic median rate in wild-type cells, consistent with silencing (Fig. 3A,C). The X element methylation rate is unaffected by loss of Sir1 (Fig. 3A,C), consistent with the observation that Sir1 is not important for phenotypic telomeric silencing (37). On the other hand, loss of Sir2, Sir3 or Sir4 results in a substantial increase in methylation rate at X elements (Fig. 3A,C). However, cluster analysis of the methylation rates of individual X elements reveals a more complex picture (Fig. S4). Four clusters were identified (Fig. S4A). X elements in cluster I are methylated at similar rates in wild type and in all four *sirΔ* mutants. X elements in cluster II are methylated faster in *sir2Δ*, *sir3Δ* and *sir4Δ* cells than in wild type and *sir1Δ* cells. In clusters III and IV, they are methylated even faster in *sir2Δ*, *sir3Δ* and *sir4Δ* cells relative to wild type and *sir1Δ* cells (Fig. S4B). The source of this variation in methylation rate is unclear, although the X elements methylated at similar rates to wild type (cluster I) tend to be the farthest from the chromosome end (> 5 kb) (Fig. S4C), whereas X element sequence similarity does not seem to be important (Fig. S4D). In conclusion, Sir2, Sir3 and Sir4 impede access to DNA at some but not all X elements.

**Fig. 3.**
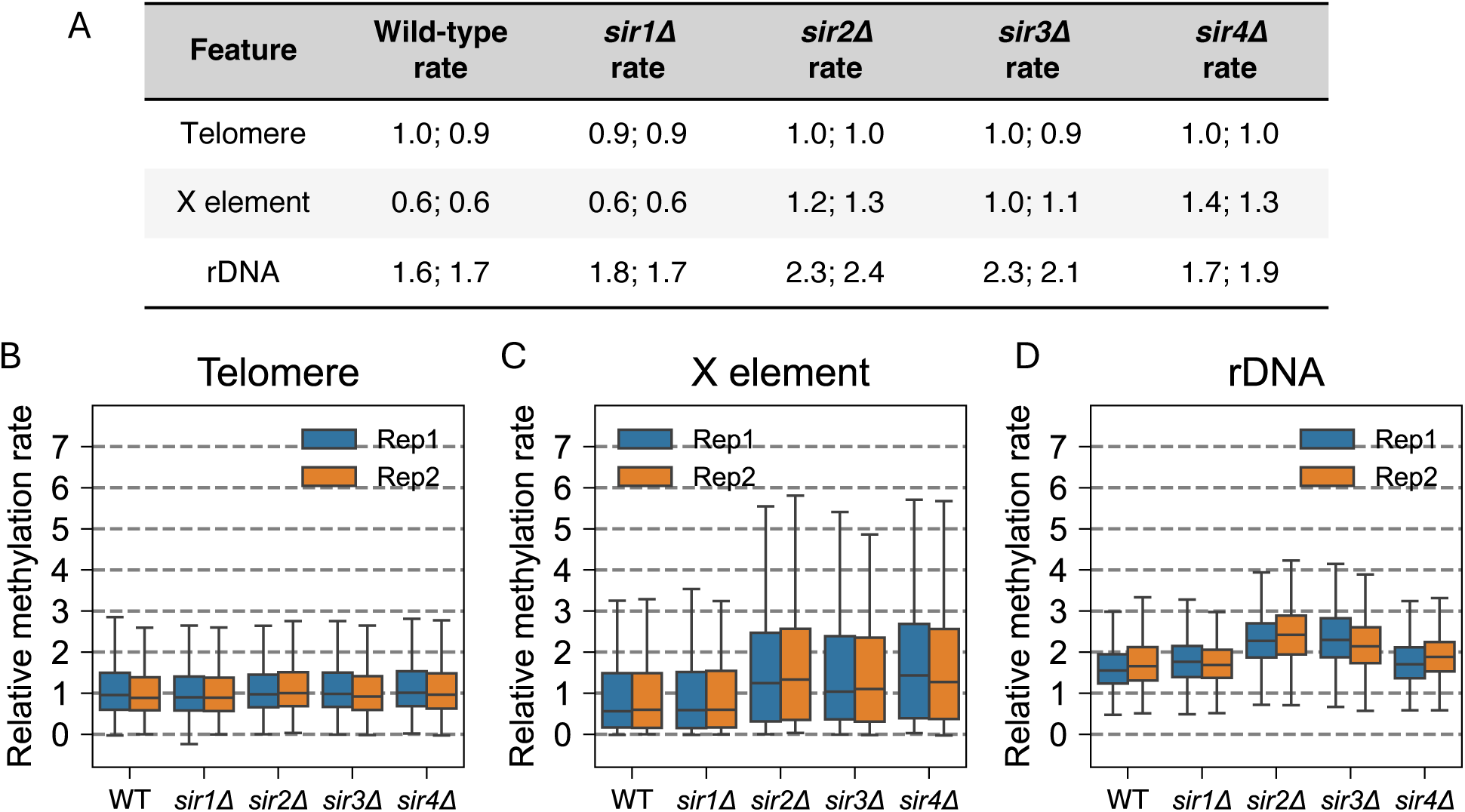
Sir proteins impede methylation of the X elements and the rDNA in vivo. (*A*) Median methylation rate constants for telomere, X element and rDNA features normalized to the whole-genome median for wild-type and *sirΔ* strains. Data from two biological replicate experiments are presented. (*B-D*) Box plots showing the distribution of individual CG methylation rate constants: (*B*) telomeres, (*C*) X elements and (*D*) rDNA (all CG sites in the *RDN37* and *RDN5* gene bodies). Boxes contain 25 to 75% of the data, the line is the median, and the whiskers represent 1.5 times the interquartile range to the farthest data points.

### Sir2 and Sir3 impede methylation of the rDNA repeats

The rDNA repeats in *S. cerevisiae* occur in a single cluster of 150 to 200 copies of a 9.1 kb sequence located on chromosome XII. Each repeat is transcribed by Pol I from a single promoter to produce a 35S transcript that is processed to yield the 5.8S, 25S and 18S rRNAs. Every repeat also contains a 5S rRNA gene transcribed by Pol III as well as coding and non-coding RNAs transcribed by Pol II. Active repeats are heavily nucleosome-depleted, whereas inactive repeats are nucleosomal; about half of the repeats are active in log phase cells (12–14).

In wild-type cells, the rDNA repeats are methylated 1.6 (replicate: 1.7) times faster than the genomic median (Fig. 3A,D). A potential explanation is that the active rDNA repeats are methylated faster than the inactive repeats because they are nucleosome-depleted, resulting in a faster methylation rate than the genomic median. The rDNA methylation rate in *sir1Δ* cells is very similar to wild type, indicating that Sir1 does not affect rDNA accessibility. In contrast, in *sir2Δ* and *sir3Δ* cells, rDNA is methylated faster than in wild-type cells, at 2.1 to 2.4 times the genomic median (Fig. 3A,D). Loss of Sir4 has little or no effect on rDNA methylation rate (Fig. 3A,D). This observation suggests that both Sir2 and Sir3 impede access to rDNA, but Sir4 does not.

### Adjacent rDNA repeats tend be methylated at similar rates in the absence of Sir2 or Sir3

More evidence that rDNA repeats adopt one of two chromatin states has come from nanopore sequencing of adjacent rDNA repeats after in vitro methylation of wild-type nuclei (38). This study revealed highly and lowly methylated populations of *RDN37* genes, presumably corresponding to active and inactive repeats, respectively, with mostly low methylation in the intergenic region, except for the Pol I promoter.

To determine the contributions of the Sir proteins to the rDNA accessibility of adjacent repeats in living cells, we analyzed our nanopore reads containing two complete rDNA repeats from the last time point (240 min). We compared the percentage methylation in adjacent *RDN37* genes in individual reads from wild-type and *sirΔ* cells by plotting the percentage methylation of *RDN37* in the lefthand rDNA repeat against the percentage methylation of *RDN37* in the righthand rDNA repeat from the same read (Fig. 4A). Pearson correlations for the wild type replicates (0.47 and 0.57) and *sir1Δ* (0.54 and 0.53) indicate that the methylation status of neighboring rDNA repeats is moderately correlated. The correlation for *sir4Δ* cells is stronger (0.64 and 0.69). The correlations for *sir2Δ* cells (0.82 and 0.81) and *sir3Δ* cells (0.76 and 0.71) are even higher. We confirmed the significance of these correlations using permutation analysis (Fig. S5).

**Fig. 4.**
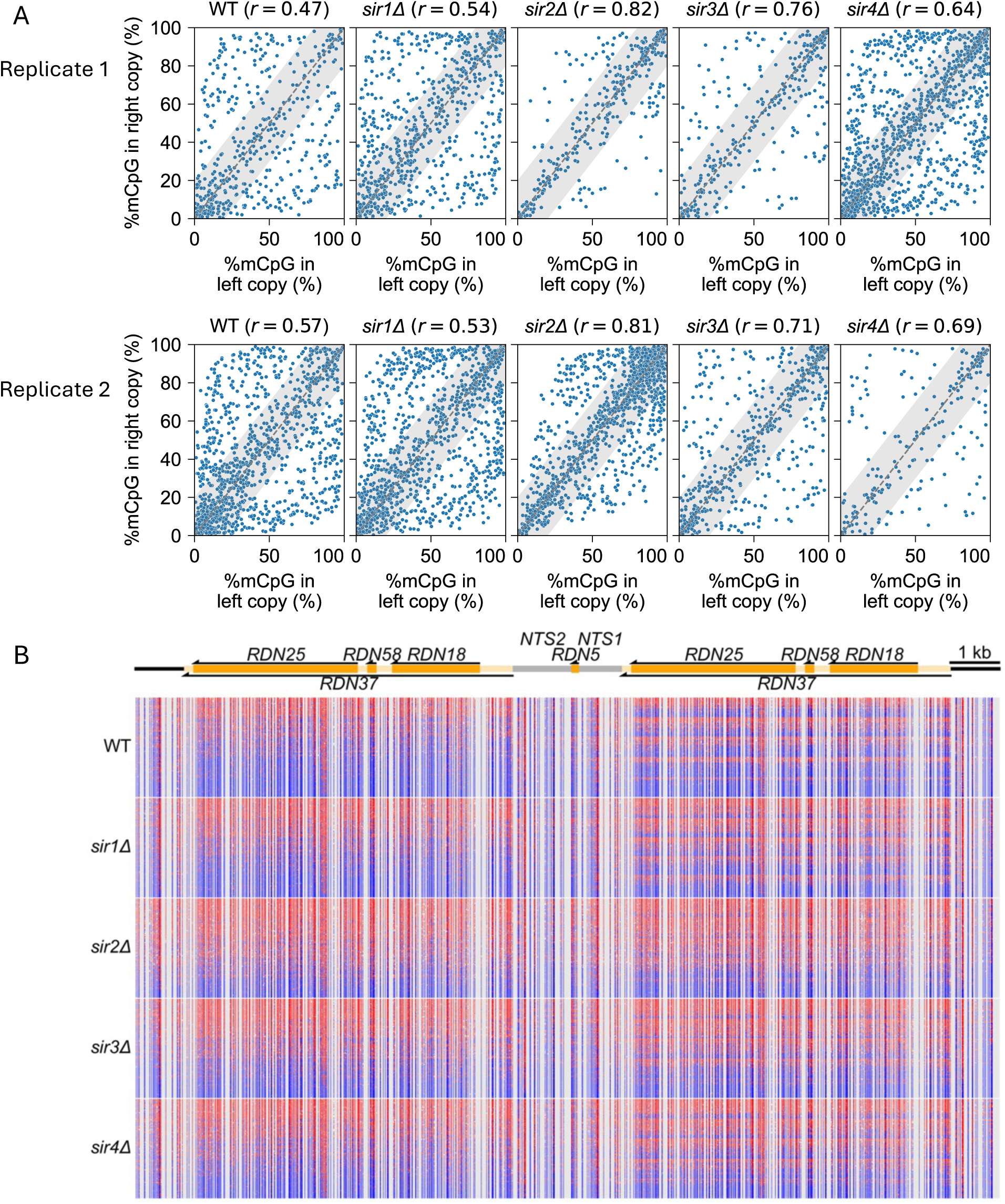
Adjacent rDNA repeats tend be methylated at similar rates in the absence of Sir2 or Sir3. Nanopore data for wild-type, *sir1Δ*, *sir2Δ*, *sir3Δ* and *sir4Δ* cells at 240 min after M.SssI induction. (*A*) Scatter plots of the fraction of methylated CG in adjacent *RDN37* genes in the same nanopore read, with Pearson correlations. The grey area indicates data within 20% of the diagonal. Differences in the number of points reflect differences in the number of reads; each sample has > 200 reads. See Fig. S5 for permutation controls. (*B*) Visualization of 200 nanopore reads containing at least two rDNA repeat elements. Reads were sorted into quintiles based on the methylated fraction of the lefthand copy of *RDN37* and sorted again within each quintile based on the methylation of the righthand copy.

To visualize these correlations, we randomly selected 200 of the nanopore reads analyzed in Fig. 4A for each strain (Fig. 4B). These reads were sorted into quintiles according to the fraction methylated of each read in the lefthand *RDN37* gene. The reads within each quintile were then sorted according to the fraction methylated in the righthand *RDN37* gene. In Fig. 4B, red indicates methylated sites, while blue indicates unmethylated sites. A read would be red on both sides if both *RDN37* genes have a high fraction of 5mCG and blue on both sides if they both have a low fraction of 5mCG. Uncorrelated repeats will appear red on one side and blue on the other, resulting in discrete striping patterns on the righthand side. Conversely, correlated repeats will result in blurred striping patterns on the righthand side. In wild-type cells, discrete stripes are apparent for the righthand repeat.

Thus, all three possibilities occur in wild-type cells: both repeats highly methylated, both lowly methylated, and one high, one low. Overall, there are fewer highly methylated rDNA repeats than lowly methylated repeats. Similar striping patterns are seen in *sir1Δ* and *sir4Δ* cells, although the fraction of highly methylated repeats is higher than in wild type. However, in *sir2Δ* cells and, to a lesser extent, in *sir3Δ* cells, the stripes are blurred, with more highly methylated repeats, and many fewer reads with one highly methylated and one lowly methylated repeat.

In summary, these data indicate that adjacent rDNA repeats in the same cell are more likely to be methylated at similar rates in the absence of Sir2 or Sir3, but not in the absence of Sir1 or Sir4. In wild-type cells, adjacent repeats are sometimes methylated at quite different rates, consistent with a difference in transcriptional activity.

## Discussion

Since nucleosomes protect their DNA from methylation in isolated nuclei, but do not do so in living cells, we proposed previously that nucleosomes must be highly dynamic in vivo, unlike in nuclei and in vitro (24, 25, 39). We suggested three models for nucleosome dynamics, based on the known properties of the ATP-dependent chromatin remodelers: removal and replacement of histone octamers, nucleosome sliding, and nucleosome conformational changes. In each case, nucleosomal DNA is rendered transiently vulnerable to methylation. We identified two genomic regions that are relatively protected from methylation in vivo: the centromeres, which are occupied by stable centromeric nucleosomes and almost completely unmethylated, and the silenced mating type loci, which are methylated much more slowly than the rest of the genome. Here we have confirmed these observations and demonstrated that slow methylation at the silenced mating type loci is dependent on Sir2, Sir3 and Sir4. Slow methylation at silenced loci suggests that Sir2, Sir3 and Sir4 suppress nucleosome dynamics, but do not completely prevent them. This effect may be direct or indirect.

Three models have been offered for transcriptional silencing of the mating loci, in which (1) Sir-associated chromatin blocks the binding of sequence-specific transcription factors (the classical steric occlusion model) (20, 40), (2) blocks pre-initiation complex formation (the pre-initiation inhibition model) (4), or (3) blocks downstream events, such as re-initiation, blocking certain elongation factors, or limiting transcriptional burst duration (5, 41).

In Fig. 5, we correlate high-resolution ChIP-exo data (42) for Sir2, Sir3 and Sir4 with our methylation rate data at the mating type loci. Sir2, Sir3 and Sir4 exhibit discrete peaks at the *HM* E and I silencers, with limited spreading between the peaks (42, 43). Within *HMLα,* there is an additional major Sir peak over the divergent *α1/α2* promoter. Adjacent to *HMRa*, there is another major Sir peak over a tRNA gene, which acts as a boundary element to prevent the spread of silencing (44). If Sir protein binding blocks access to the DNA, we would expect that the Sir protein peaks would correlate with the slowest methylation rates. However, the opposite is true at five of the six major Sir peaks in the mating loci. At *HMRa*, methylation is faster within the Sir peaks at the E and I silencers and the tRNA gene than at sites in the *a1* gene, the *a2* gene and the *Ty1_HMRa* transposable element (Fig. 2A; Fig. 5). At *HMLα*, methylation is faster at the I silencer and the *α1/α2* promoter, but slow at the E silencer and the *α1*and *α2* genes (Fig. 5). Faster methylation at the Sir peaks could be explained by the fact that all six Sir peaks coincide with NDRs (21, 22) (Fig. 5), but this does not account for slow methylation at the *HMLα* E silencer.

**Fig. 5.**
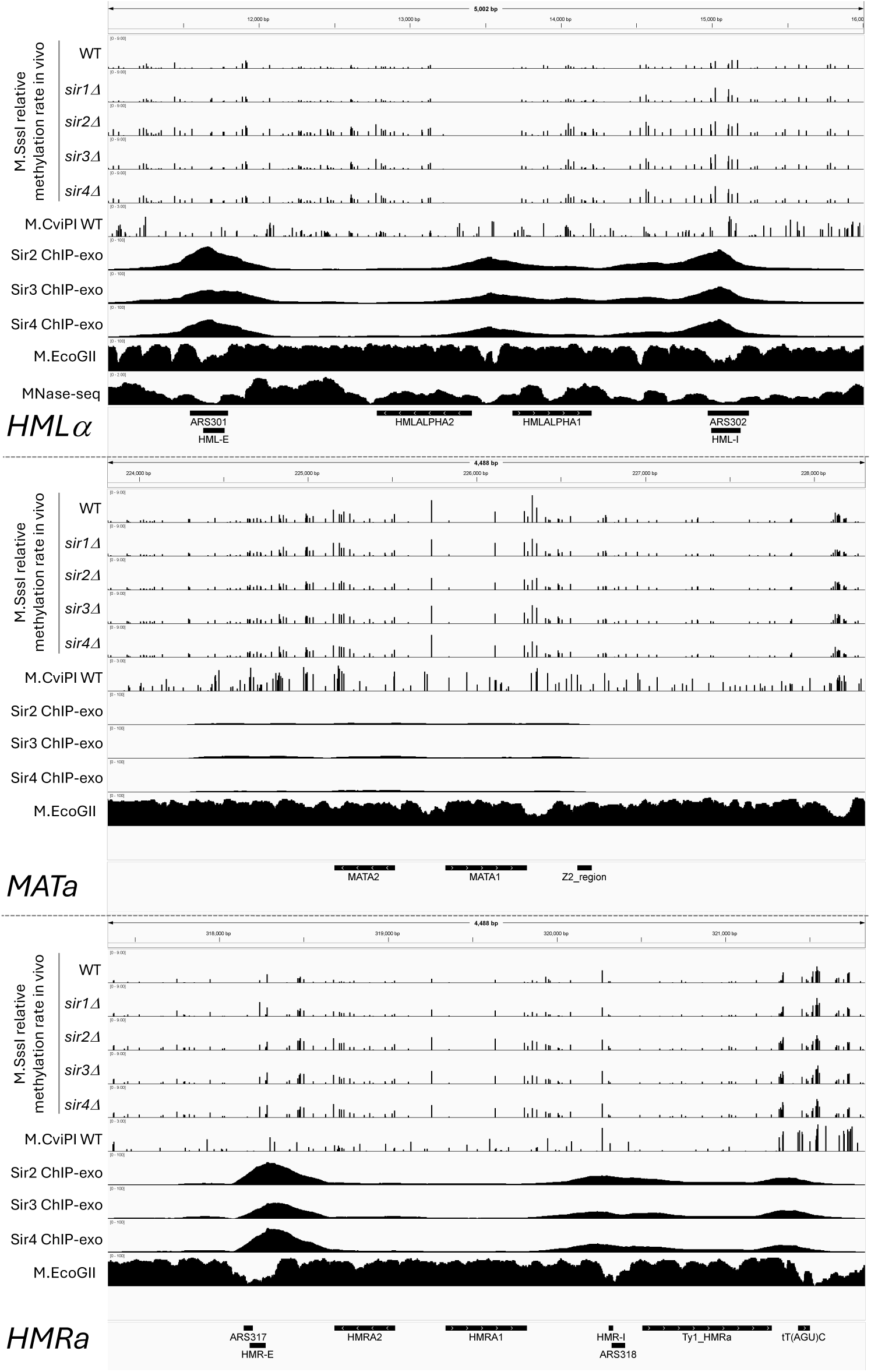
Methylation rates at CG sites in *HMLα*, *MATa* and *HMRa* relative to Sir protein location. IGV tracks. M.SssI methylation rate data for wild-type (WT), *sir1Δ*, *sir2Δ*, *sir3Δ* and *sir4Δ* strains, and wild-type data for M.CviPI. M.EcoGII data (57) and MNase-seq data (29) for wild-type nuclei. ChIP-exo data for the Sir proteins (42).

The Sir proteins also attenuate methylation rates at most of the telomeric X elements. ChIP-exo data for Sir2, Sir3 and Sir4 (42) indicate a major peak over the X element with some spread to cover the entire element and some of the DNA between the X element and the chromosome end (42) (Fig. 6A). In wild-type cells, methylation is very slow in the main Sir peak, but faster on each side of the main peak. In the absence of Sir2, Sir3 or Sir4, the methylation rate increases. These rate differences might be explained by the well-positioned nucleosome that is coincident with the main Sir peak (10, 42), which might have particularly slow dynamics.

**Fig. 6.**
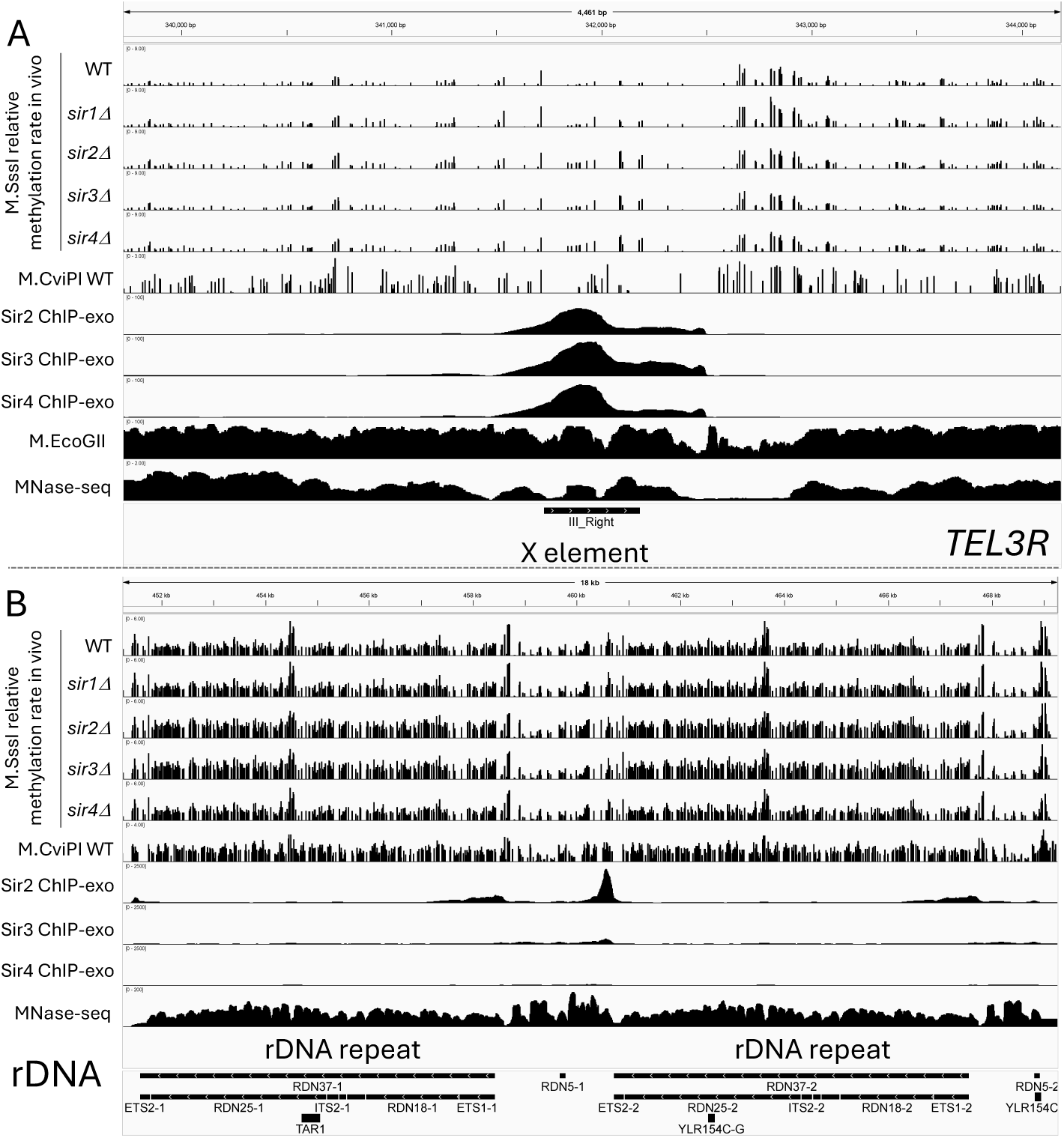
Methylation rates at a representative telomere and at rDNA relative to Sir protein location. IGV tracks. M.SssI methylation rate data for wild-type (WT), *sir1Δ*, *sir2Δ*, *sir3Δ* and *sir4Δ* strains, together with wild-type data for M.CviPI. M.EcoGII data (57). and MNase-seq data (29) for wild-type nuclei. ChIP-exo data for the Sir proteins (42). (*A*) *TEL3R* (the telomere at the righthand end of chromosome III) including the X element. (*B*) Two rDNA repeats (chromosome XII).

At the rDNA locus, wild-type methylation rates are high relative to the rest of the genome, likely because of the nucleosome-depleted active rDNA repeats that are methylated more quickly (12–14). Thus, the observed median methylation rate for the rDNA will depend on the fraction of active repeats. Overall, in the absence of Sir2 or Sir3, the median rate is faster than in wild-type cells. Moreover, our analysis of adjacent *RDN37* repeats in wild-type cells indicates that adjacent repeats are frequently methylated at quite different rates. In cells lacking Sir2 or Sir3, adjacent repeats are more likely to be methylated at similar rates. This observation may indicate that the Sir proteins silence rDNA repeats randomly, resulting in a weaker correlation between the methylation rates of adjacent repeats.

ChIP-exo data (42) indicate a major Sir2 peak over the non-transcribed region 3’ of *RDN37* (*NTS1*) and a minor Sir2 peak over the Pol I promoter (*NTS2*), with some limited spreading of Sir2 (Fig. 6B). Sir3 and possibly Sir4 (the Sir4 signal is very weak) spread in between these two Sir2 peaks to cover the entire intergenic region, including the 5S rRNA gene transcribed by Pol III. Unlike the Sir2 peaks at the silenced mating type loci and the X elements, the Sir2 peaks at the rDNA are not coincident with the Sir3 and Sir4 peaks. The Sir protein distribution at the rDNA is more consistent with the classical spreading model (45), although there is virtually no Sir protein spreading over the *RDN37* gene body (Fig. S6). These data suggest that Sir protein binding in the spacer DNA between *RDN37* genes is sufficient for rDNA silencing.

We observed a relatively minor increase in methylation rate in the *sir1Δ* mutant at *HMRa* but not at *HMLα*. Previous studies on silencing, ranging from classical mating assays (34) to more modern fluorescence reporter assays (33, 35), observed that the absence of Sir1 results in two phenotypically distinct cell populations: one where silencing is maintained, and one where it is not. Therefore, it was proposed that Sir1 is primarily involved in the de novo establishment of silencing once the state has been lost. Recent studies also provide some evidence for a role in maintenance: a temporally sensitive Cre recombinase assay captured increased frequency of transient loss-of-silencing events in the absence of Sir1 (35). Moreover, when artificially recruited in a dose-dependent manner, Sir1 can attenuate the expression of genes that are strong enough to overwhelm transcriptional repression at a silent locus (5). We observe that the methylation rate at *HMLα* is unaffected in *sir1Δ* cells, whereas *HMRa* is methylated somewhat faster than in wild type cells. This is consistent with the fact that *HMRa* is de-repressed in a higher proportion of *sir1Δ* cells than *HMLα* (33, 35). However, we note that we are measuring relative accessibility, which may not relate directly to expression data (33).

In summary, it seems unlikely that steric occlusion is the primary mechanism through which transcriptional silencing is achieved, since silenced DNA is still accessible to methyltransferases in living cells, albeit at a slower rate than elsewhere in the genome. Sir protein mobility in vivo could explain how Sir proteins impede, but not prevent, access to silenced DNA (3, 46). Other potential contributing factors to transcriptional silencing (3) include: (1) Sir2-mediated histone deacetylation increasing the affinity of histone tail domains for DNA; (2) inhibition of an ATP-dependent remodeling activity by histone deacetylation; and (3) putative formation of Sir-mediated condensed chromatin loops. Such loops might involve local interactions between elements defined by Sir2 peaks, and/or long-range Sir-dependent interactions between *HMLα* and *HMRa* and with telomeres (43, 47–49). We propose that the Sir proteins define chromatin domains in which nucleosome dynamics are reduced, but not eliminated, resulting in slower methylation than elsewhere in the genome.

## Materials and Methods

### Plasmids

Plasmids p923, p924, and p925 were synthesized by Thermo Fisher GeneArt to contain integration cassettes with homology arms targeting *SIR2*, *SIR3* and *SIR4*, respectively, to replace the coding region with the hygromycin resistance gene (*Hph*) in the opposite orientation. The integration plasmid targeting *SIR1* (p994) was constructed by inserting a similarly oriented NatNT2 gene into p984 (Thermo Fisher), which contains *SIR1* homology arms. The integration plasmid (p956) for Gcn4-dependent expression of codon-optimized M.CviPI with an SV40 nuclear localization signal, an auxin-dependent degron and three HA tags fused to its N-terminus was constructed as follows: A synthetic plasmid (p919; Thermo Fisher GeneArt) containing a 1479-bp AvrII-AgeI insert in which the truncated, modified *tCUP1* promoter (the proximal UAS was replaced with a high-affinity Gcn4 binding site as described previously (24) is fused to the M.CviPI open reading frame (ORF) (with optimal yeast codons and potential splice sites removed) with a C-terminal degron and two HA tags. This 1479-bp AvrII-AgeI insert in p919 was used to replace the 1227-bp AvrII-AgeI Dam fragment in p876 (24) to yield p921, an integration plasmid for SM-induced expression from the *tCUP1* promoter of M.CviPI with a C-terminal degron and 3 HA tags. However, this enzyme was inactive in vivo, even though it was expressed, presumably due to the C-terminal tag. Consequently, another synthetic plasmid was obtained (p954; Thermo Fisher GeneArt) containing a 1542-bp AvrII-NheI insert, in which an SV40 nuclear localization signal, a degron and 3 HA tags were fused to the N-terminus of the codon-optimized M.CviPI ORF. This 1542-bp fragment was used to replace the 1509-bp AvrII-NheI fragment in p921 to create p956. A NotI-EcoRV digest of p956 was used for integration at the *HO* locus (the digest was only partial, apparently due to leaky expression of M.CviPI in *E. coli*, resulting in some methylation of the NotI sites, blocking NotI). All plasmids were confirmed by nanopore sequencing and are available upon request.

### Strains

The yeast strains used in this study are listed in Table S1. For *sir2Δ* (YPE840), *sir3Δ* (YPE841) and *sir4Δ* (YPE842) strains, YHP827 (24) was transformed with NotI digests of p923, p924 or p925, respectively, and selecting on yeast peptone dextrose (YPD) plates with hygromycin. The *sir1Δ* strain (YKW874) was constructed by transforming YHP827 with a NotI digest of p994 and selection for nourseothricin resistance on YPD plates. YHP853 was obtained by transforming YDC111 (50) with a NotI/EcoRV digest of p956, followed by selection for G418 resistance.

### M.SssI and M.CviP1 methylation assays

YHP827 and its derivatives YPE840, YPE841, YPE842 and YKW874 were inoculated in 250 mL 2% dextrose synthetic complete medium without isoleucine and valine (SC -ile -val) and grown at 30°C to OD_600_ of 0.6 – 0.8. For the first time point immediately before M.SssI induction (0 min), 50 mL culture was taken for the sample, of which 40 mL was reserved for DNA extraction and 10 mL for protein extraction. Cell pellets were stored at -70°C. To induce M.SssI expression, SC -ile -val was replenished to 225 mL and supplemented with SM (Sigma-Aldrich 324224, 2 mg/mL in DMSO) to a final concentration of 1 µg/mL. Cell pellets for time points after M.SssI induction (30, 60, 120 and 240 min) were obtained and stored as per the first time point. YHP853 cells expressing M.CviPI were cultured and processed as described above, except that an extra time point, at 480 min, was taken.

### DNA extraction

Cell pellets were permeabilized and washed twice with 500 µL of 5x TE (50 mM Tris-HCl pH 8.0, 5 mM EDTA) + 2% SDS. Next, to remove the SDS, they were washed twice more with 500 µL 5× TE, then resuspended in 450 µL 5x TE + 15 mM 2-mercaptoethanol. To digest cell walls, 50 µL lyticase (Sigma-Aldrich L2524, 25,000 units/mL) was added, mixed gently by inversion, then incubated at 37°C for 10 min. To stop the reaction, 50 µL 20% SDS was added, mixed gently, and incubated at room temperature for 5 min. DNA was isolated by addition of 110 µL 5 M potassium acetate, followed by two 1x volume chloroform extractions, precipitating and pelleting with 0.7x volume isopropanol, and washing with 70% ethanol. The resulting pellet was air-dried and dissolved in 100 µL 10 mM Tris-HCl pH 8.0, 0.1 mM Na-EDTA, 0.4 mg/mL RNase A and incubated at 37°C for 2 h. Further purification of DNA was performed with the PureLink Genomic DNA Mini Kit (Invitrogen K182002). 10 µL Proteinase K and 10 µL RNase A, supplied with the kit, were added to each sample and incubated at room temperature for 2 min. 100 µL PureLink Genomic Lysis/Binding Buffer was added to each sample, mixed gently by inversion, and incubated at 55°C for 10 min. Finally, 100 µL ethanol was added to each sample and mixed gently by inversion before proceeding to the “Binding DNA” step of the manufacturer-provided protocol and completing the extraction as per instructions.

### Nanopore sequencing

Genomic DNA samples were barcoded with the Native Barcoding Kit (SQK-NBD114.24) from Oxford Nanopore Technologies (ONT) as per manufacturer instructions and sequenced with R10.4.1 flow cells on a MinION Mk1C instrument running MinKNOW 24.11.8 (ONT). Reads were base-called using Dorado v0.9.1 with the dna_r10.4.1_e8.2_400bps_sup v5.0.0 base-calling model and 5mCG_5hmCG v3 modification model (--min-qscore 8). Reads were aligned to sacCer3 or the YHP827 genome (24) using the Dorado aligner. For rDNA repeat analysis, a synthetic repeat array was constructed by extracting the rDNA repeat unit from chromosome XII (coordinates 451575-460711) and assembling two complete repeats flanked by 2 kb partial sequences. Reads were aligned to this custom reference using the Dorado aligner. 5mCG sites were predicted using modkit v0.4.3 (--cpg --combine-strands --ignore h). For single-read methylation analysis, reads overlapping target regions were extracted and 5mCG sites in individual reads were annotated with a threshold >0.8 (unmethylated CG sites are < 0.2). Only reads containing at least one 5mCG were retained.

### Computational analysis

Genome-wide methylation rates were calculated from the median methylated fraction across all chromosomes except for mitochondrial and 2-micron plasmid DNA, assuming pseudo first-order kinetics. Each data set was normalized to its genomic median (set at 1). We annotated the E and I silencers in the *HM* loci and the X elements in the YHP827 genome based on the sacCer3 assembly using MUMmer v4.0.13 (51). We annotated the *MATa* locus as the region that is identical to the *HMRa* locus. The telomeres in the YHP827 genome are defined as the region from the X element to the chromosome end according to SGD. Published ChIP-seq data (GSE147927 (42, 52) were pre-processed by fastp/0.24.0 (53), then aligned to the sacCer3 or the YHP827 genome using Bowtie2 v2.5.3 (-X 5000 --very-sensitive --no-discordant --no-mixed --no-unal) (54). Aligned reads were filtered using SAMtools v1.21 (-f 0x2 -F 0x300 -q 1) (55). Read occupancy was calculated using BEDTools v2.31.1 (56). We calculated the average occupancy of nuclear chromosomes (excluding chrXII) and normalized the ChIP-exo seq data to the average occupancy (set to 1.0). We used MNase-seq data from GSE69400 (29). Genome accessibility data from M.EcoGII methyltransferase mapping in nuclei (57) were incorporated for comparative analysis. Data visualization was performed using Integrative Genomics Viewer (IGV)(58). We developed Python scripts to analyze the data. This software used the following packages: h5py, Matplotlib (59), NumPy (60), pandas (61), pyBigWig, pysam, SciPy (62), seaborn (63) and statsmodels (64).

### SDS PAGE and Western blotting

Cell pellets were resuspended in NuPAGE lithium dodecyl sulfate buffer (Invitrogen NP0007) with 20 mM 2-mercaptoethanol (5 OD_600_ units of cells per 100 µL) and heated at 99°C for 5 min. Samples were centrifuged at 9,400 *g*; the supernatant was stored at -20°C. For SDS PAGE, samples were thawed and heated at 70°C for 10 min before loading into two 4-12% Bis-Tris NuPAGE Midi Gels (Invitrogen WG1403BOX) in an XCell4 Surelock Midi-Cell (Invitrogen WR0100) and run with NuPAGE MOPS SDS Running Buffer (Invitrogen NP0001) as per manufacturer instructions. One gel was used for Coomassie staining; the other gel was transferred to a PVDF membrane with the iBlot Dry Blotting System (Invitrogen IB1001) with regular sized transfer stacks (Invitrogen IB401001) per manufacturer instructions. To detect HA, the membrane was placed in a hybridization tube and blocked with 20 mL 5% skim milk in TBST (20 mM Tris-HCl pH 8.0, 0.5 M NaCl, 0.1% Tween 20) with rotation at room temperature for 1 h, followed by hybridization in 10 mL 1:5,000 dilution of an HRP-conjugated α-HA antibody (Roche F310) in 5% skim milk TBST blocking buffer with rotation at 4°C overnight. The membrane was washed three times with 20 mL TBST with rotation at room temperature for 10 min, then incubated with SuperSignal West Pico PLUS Chemiluminiscent Substrate (Thermo Scientific 34577) using 2 mL of a 1:1 mixture of the substrate and stable peroxide components from the kit at room temperature for 5 min to develop. The membrane was imaged for HA using the Azure 300 Chemiluminescent Western Blot Imager (Azure Biosystems AZI300-01). To detect tubulin, the membrane was returned to a hybridization tube and stripped with 20 mL PBST (phosphate buffered saline, 0.1% Tween 20) with rotation at room temperature for 1 h and blocked with 20 mL 5% skim milk in PBST with rotation at room temperature for 1 h. The membrane was hybridized in 5 mL 1:20,000 dilution of an HRP-conjugated α-tubulin antibody (Abcam ab185067) in 5% skim milk PBST blocking buffer with rotation at room temperature for 1 h, followed by three 15 mL PBST washes with rotation at room temperature for 10 min each. The membrane was developed and imaged for tubulin as above.

## Data availability

Nanopore sequencing data are available at the GEO database with the accession number GSE311530. Scripts are available at GitHub: https://github.com/zhuweix/Yeast_Sir_2025

## Acknowledgements

We thank Rohinton Kamakaka for very helpful comments on the manuscript. This study used the high-performance computational capabilities of the Biowulf Linux cluster at the NIH. This research was supported by the Intramural Research Program of the National Institutes of Health (NIH). The contributions of the NIH authors are considered Works of the United States Government. The findings and conclusions presented in this paper are those of the authors and do not necessarily reflect the views of the NIH or the U.S. Department of Health and Human Services.

## Author contributions

K.W., H.P. and P.E. performed the experiments; Z.X. performed the bioinformatic analysis; K.W. and D.C. wrote the manuscript.

## SI Appendix

Figures S1-S6 and Table S1.

