## Supplemental Information for "Sir proteins impede, but do not prevent, access to silent chromatin in living *Saccharomyces cerevisiae*"

Kenneth Y. Wu[1], Zhuwei Xu[1], Hemant K. Prajapati, Peter R. Eriksson and David J. Clark*
Division of Developmental Biology, *Eunice Kennedy-Shriver* National Institute of Child Health and Human Development, National Institutes of Health, Bethesda MD 20892, USA

Figures S1-S6.

Table S1.

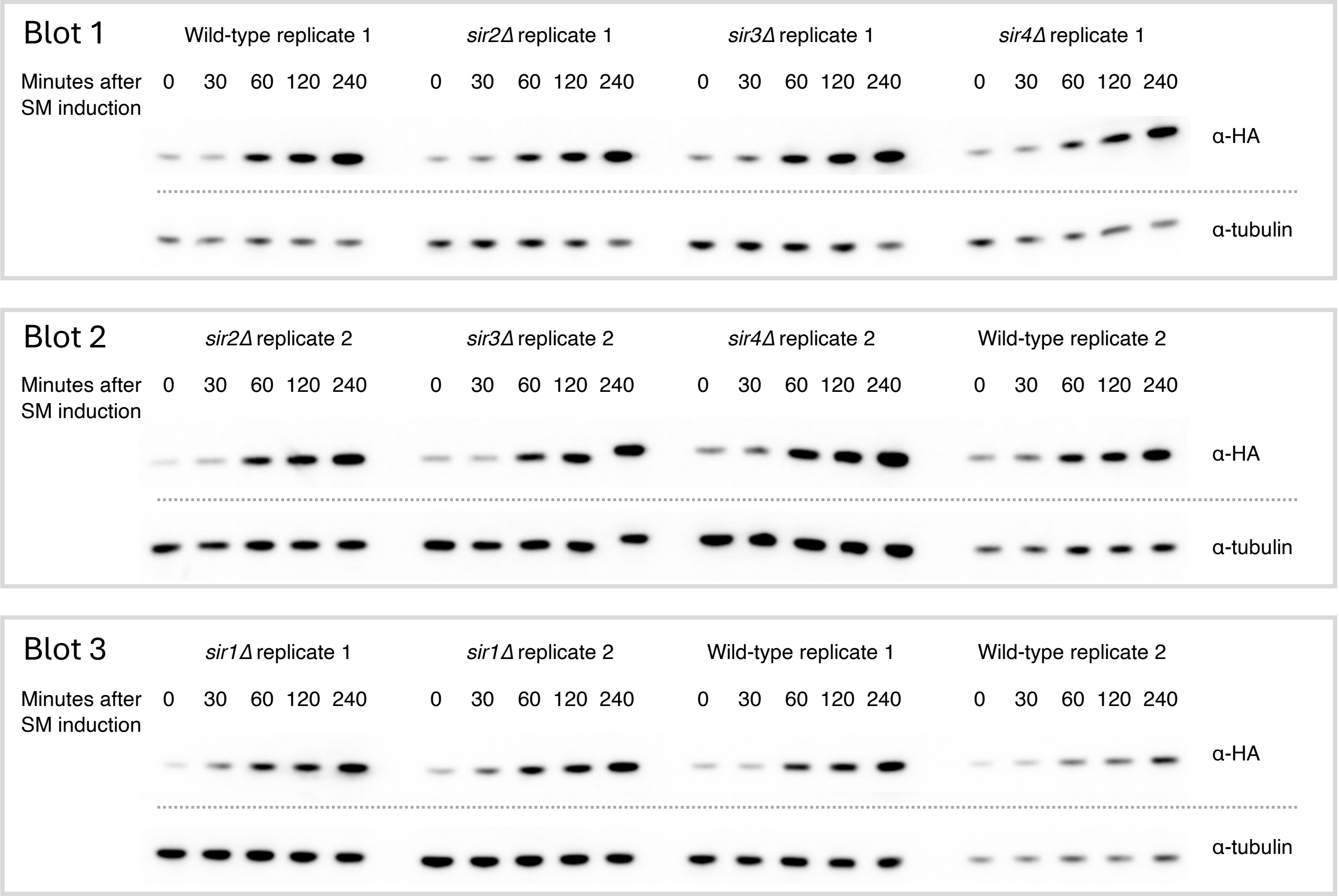


**Fig. S1**. SM induction of M.SssI expression in replicate wild-type, *sir1Δ*, *sir2Δ*, *sir3Δ* and *sir4Δ* experiments. Western blots for HA-tagged M.SssI and for tubulin across time course samples for each strain. Each blot contains four time courses, probed first for M.SssI-3HA (upper rows) and then for tubulin (lower rows).

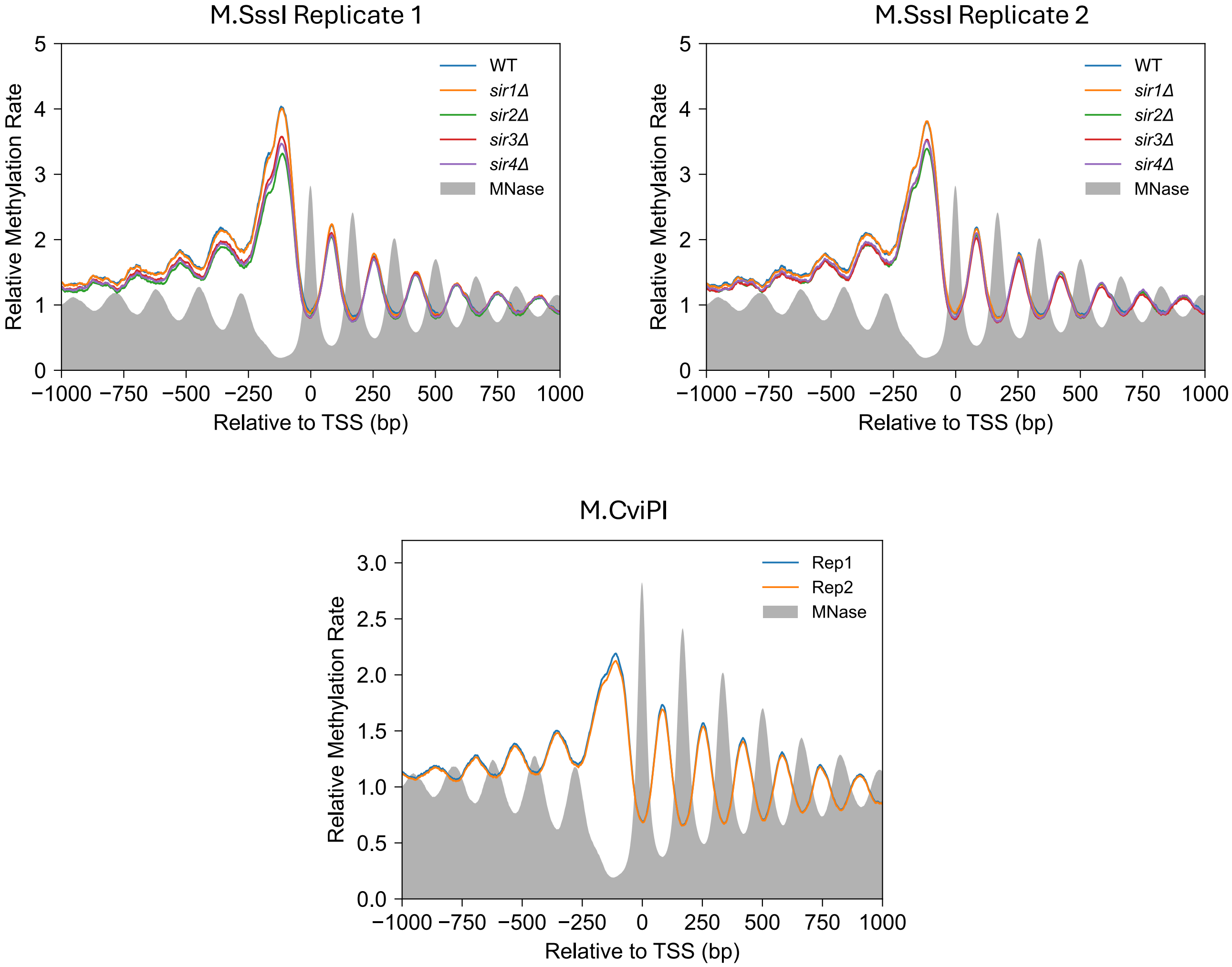


**Fig. S2**. Global nucleosomal phasing at genes is unaffected by the absence of Sir1, Sir2, Sir3 or Sir4 in living cells. Nucleosomal phasing can be detected from aggregate M.SssI methylation rates across all genes aligned to the +1 nucleosome for WT, *sir1Δ*, *sir2Δ*, *sir3Δ* and *sir4Δ* strains. Data for M.CviPI methylation rates in wild-type cells are also shown. MNase-seq data for the same regions in a wild-type strain are represented in grey.

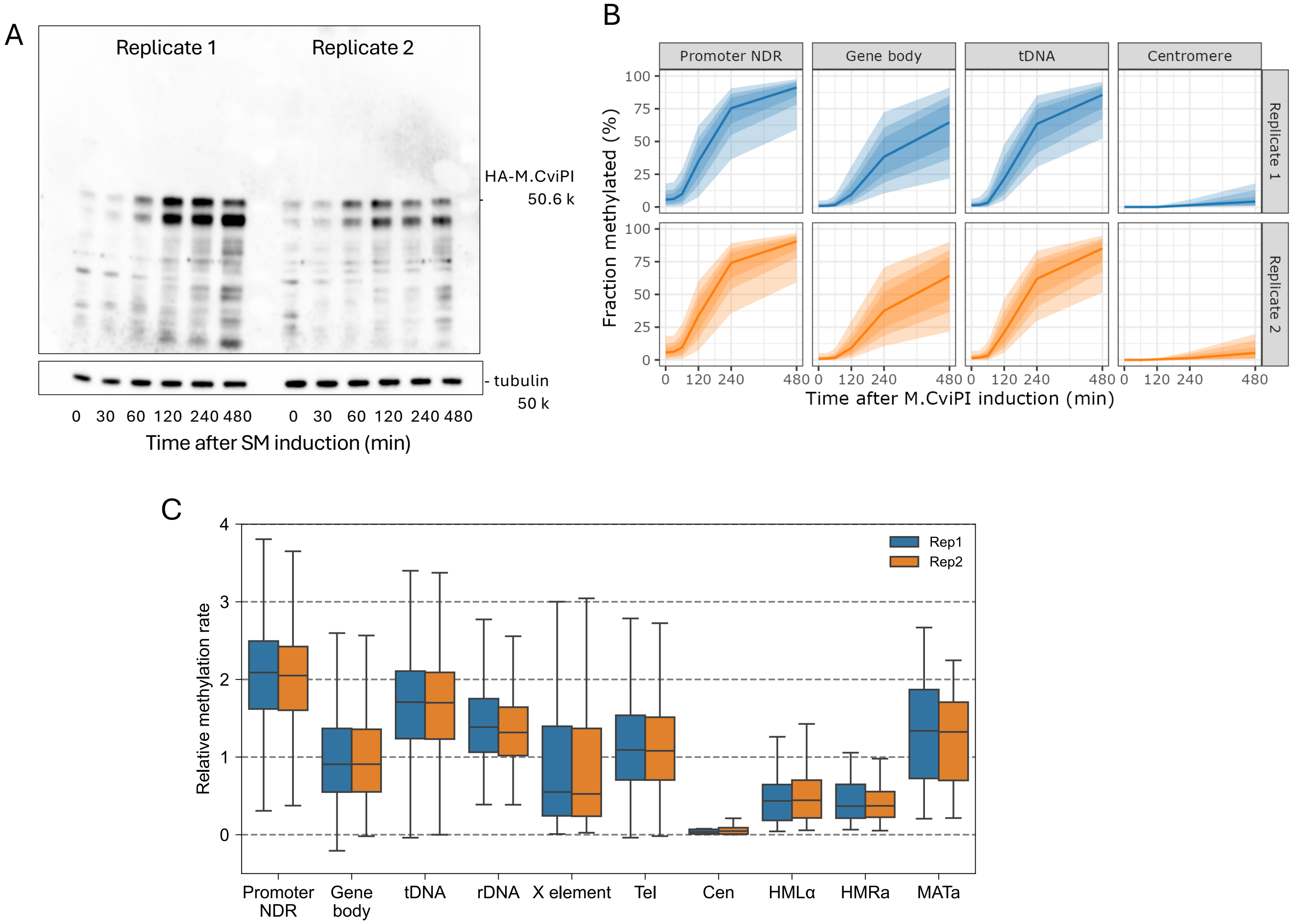


D

| Rep | Prom. NDR | Gene body | tDNA | rDNA | XE | TEL | CEN | *HMLα* | *MATa* | *HMRa* |
|---|---|---|---|---|---|---|---|---|---|---|
| 1 | 2.1 | 0.9 | 1.7 | 1.4 | 0.6 | 1.1 | 0.0 | 0.4 | 1.3 | 0.4 |
| 2 | 2.1 | 0.9 | 1.7 | 1.3 | 0.5 | 1.1 | 0.1 | 0.4 | 1.3 | 0.4 |

**Fig. S3**. M.CviPI methylation assays mirror M.SssI methylation assays. (*A*) Western blot for HA-tagged M.CviPI and tubulin. Two biological replicate experiments were compared in the same blot. (*B*) M.CviPI methylation time courses showing the median GC site methylation (solid line) for promoter NDRs, gene bodies, tDNA and centromeres. Shading: lightest to darkest: 5-95%, 15-85%, and 25-75% of all GC sites in the feature. (*C*) Box plots showing the distributions of methylation rate constants for all individual GC sites in each genomic feature. Boxes contain 25 to 75% of the data, the line is the median and the whiskers represent 1.5 times the interquartile range to the farthest data points. (*D*) Table of median methylation rate constants of GC sites for each genomic feature (XE: X elements).

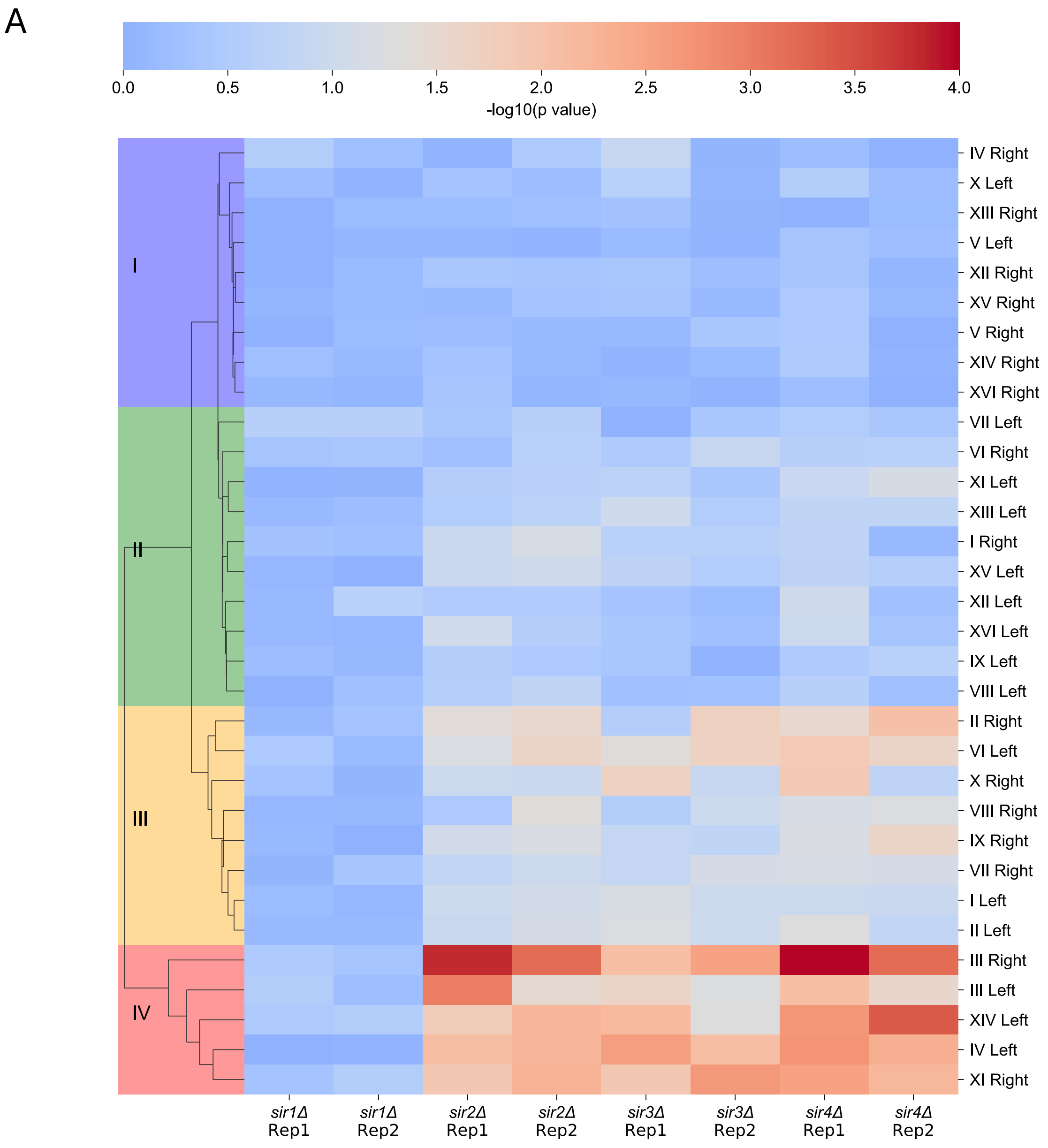


*See following pages for B, C and D.*

**Fig. S4**. X element classification and methylation rate differences. (*A*) Heatmap showing -log10(p values) from Mann-Whitney *U* tests comparing X element methylation rates of individual CpG sites for pooled WT (replicates 1 and 2) and separate *sir* mutant replicates. Each column represents the comparison between pooled WT and one biological replicate of the indicated *sir* mutant. Each row represents one X element. The colormap is centered at p = 0.05 (-log10(0.05) = 1.3), with blue indicating p > 0.05 (insignificant) and red indicating p < 0.05 (significant). We derived four clusters of X elements (I-IV). (*B – next page)* Box plot analysis of methylation rate constants for each X element (organized by the clusters in *A*). (*C – following page*) Box plots showing the distances of X elements from the chromosome end in each cluster. X elements in cluster I are all > 5 kb from the chromosome end. (*D - following page*) X element sequence identity matrix (Clustal Omega). No strong correlation between methylation rate cluster and sequence identity.

B

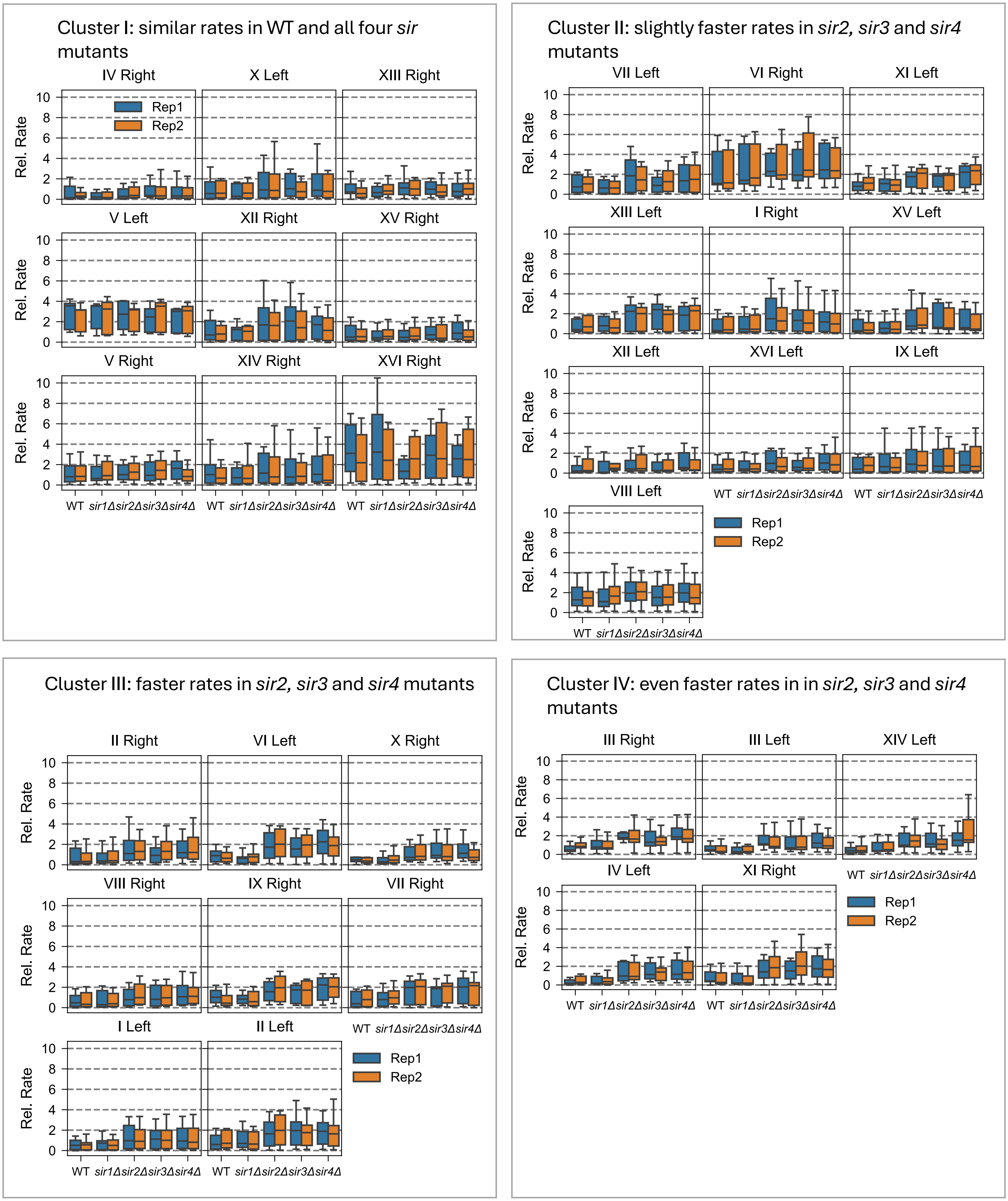


**Fig. S4**, *continued*.

C

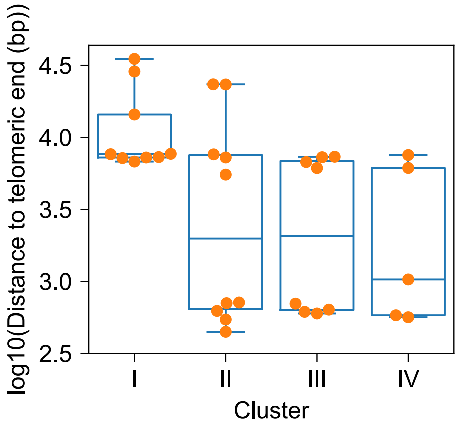


D

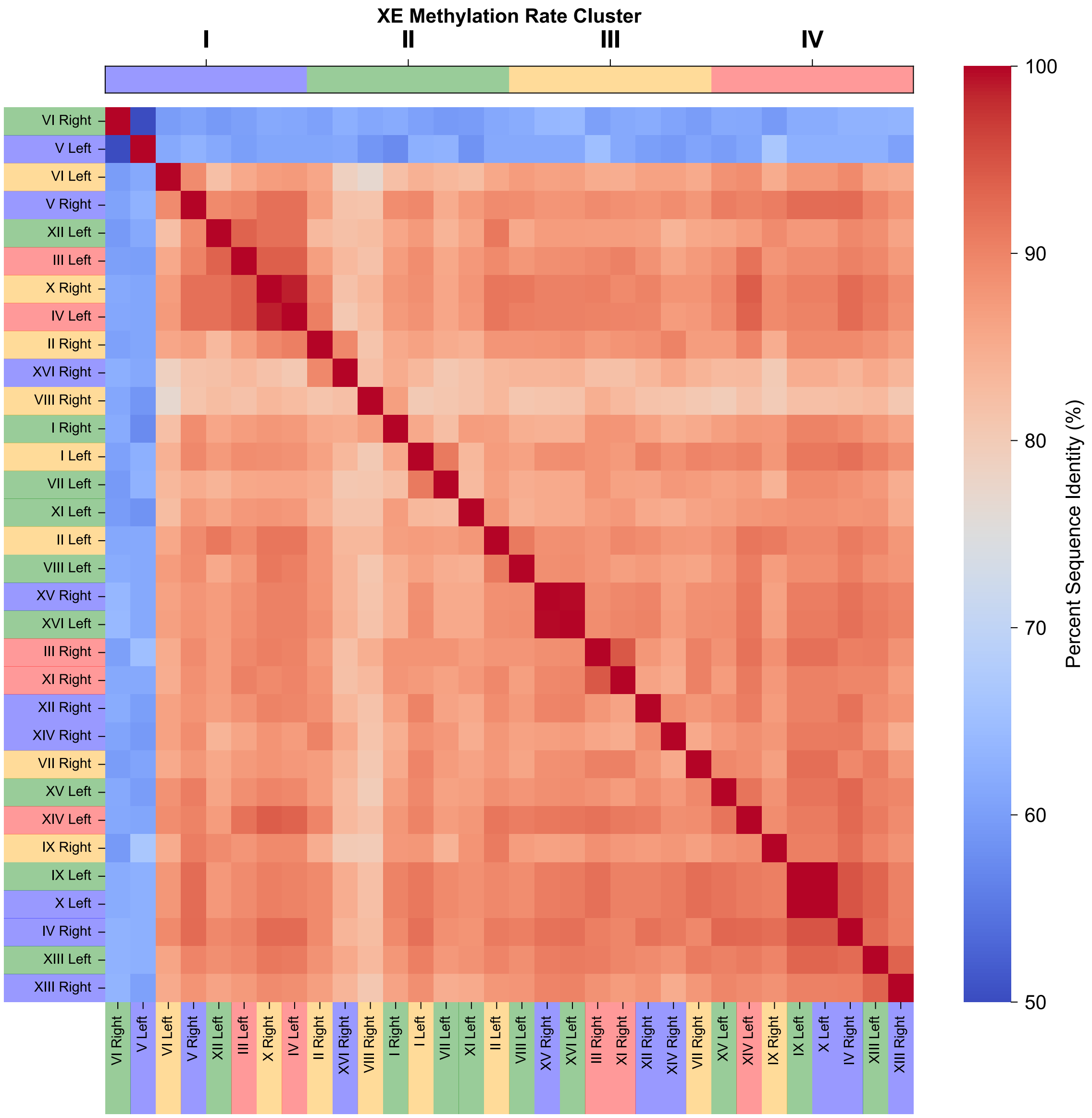


**Fig. S4**, *continued*.

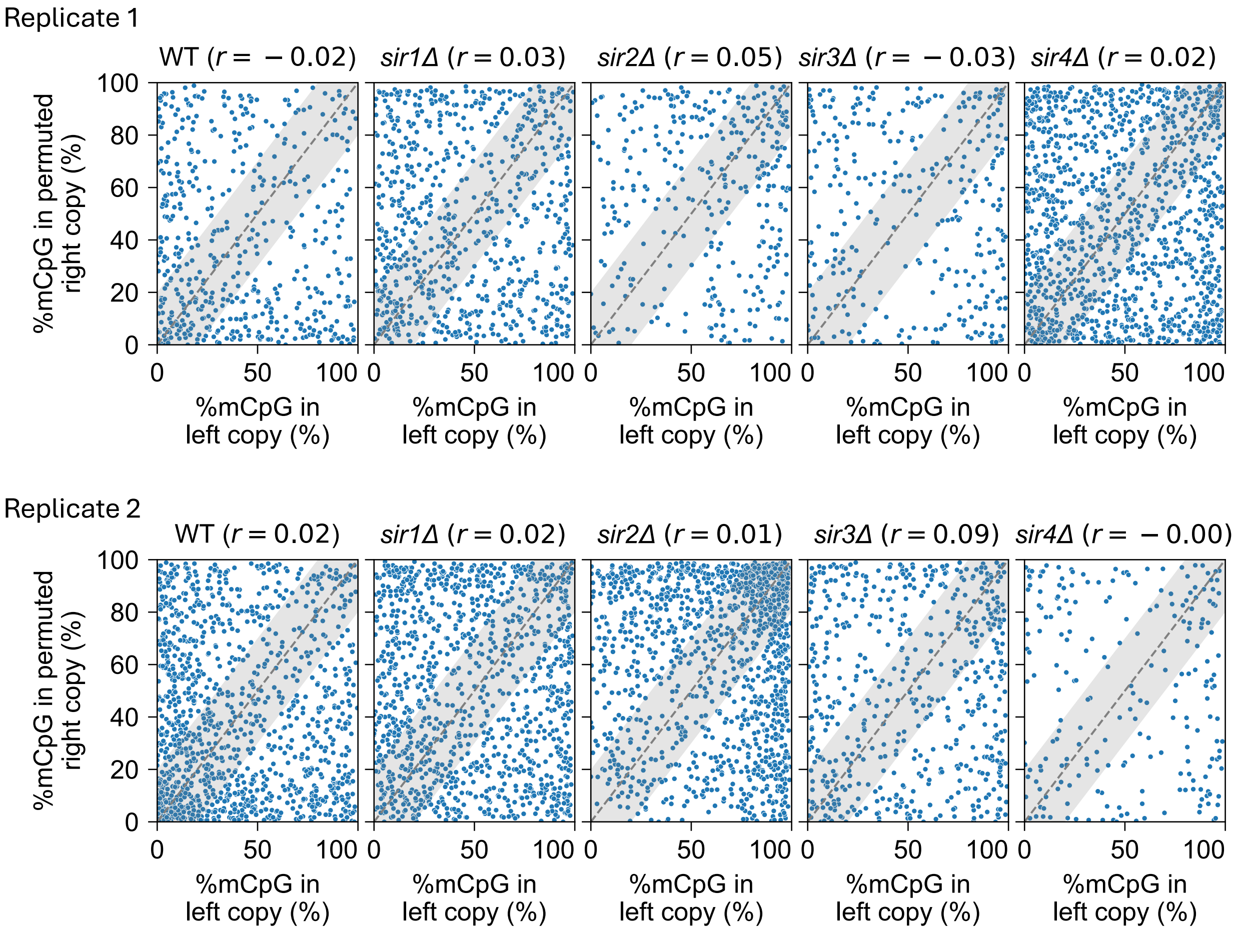


**Fig. S5.** Permutation controls for the correlation between M.SssI methylated fractions for adjacent *RDN37* genes in the same nanopore read (see Fig. 4A). Methylated fractions of the left *RDN37* copy were randomly shuffled while the methylated fractions of the right copy were unaltered. The data were then re-plotted to assess the significance of the observed left-right correlation. The grey area indicates data within 20% of the diagonal.

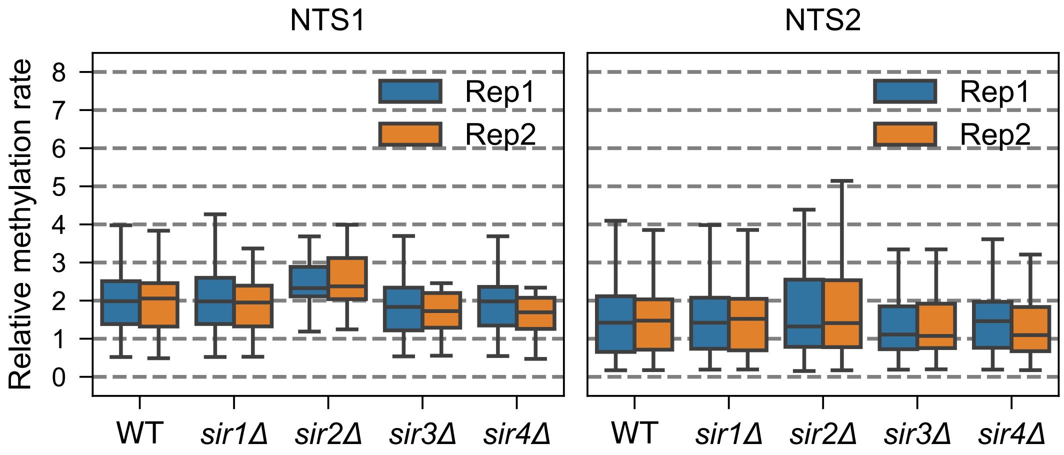


**Fig. S6**. Box plots showing the distributions of methylation rate constants of individual CG sites located within the *NTS1* and *NTS2* regions of the rDNA repeats. Boxes contain 25 to 75% of the data, the line is the median and the whiskers represent 1.5 times the interquartile range to the farthest data points.

**Table S1**. Yeast strains used in this study.

| Strain | Genotype |
|---|---|
| **YDC111** | *MATa ade2-1 can1-100 leu2-3,112 trp1-1 ura3-1* |
| **YHP827** | *MATa ade2-1 can1-100 leu2-3,112 trp1-1 ura3-1 ho::TIR1_MSssI-degron-3HA_KanMX (p906)* |
| **YPE840** | *MATa ade2-1 can1-100 leu2-3,112 trp1-1 ura3-1 ho::TIR1_MSssI-degron-3HA_KanMX (p906) sir2Δ::Hph (p923)* |
| **YPE841** | *MATa ade2-1 can1-100 leu2-3,112 trp1-1 ura3-1 ho::TIR1_MSssI-degron-3HA_KanMX (p906) sir3Δ::Hph (p924)* |
| **YPE842** | *MATa ade2-1 can1-100 leu2-3,112 trp1-1 ura3-1 ho::TIR1_MSssI-degron-3HA_KanMX (p906) sir4Δ::Hph (p925)* |
| **YHP853** | *MATa ade2-1 can1-100 leu2-3,112 trp1-1 ura3-1 ho::TIR1_NLS-degron-3HA-MCviPI_KanMX (p956)* |
| **YKW874** | *MATa ade2-1 can1-100 leu2-3,112 trp1-1 ura3-1 ho::TIR1_MSssI-degron-3HA_KanMX (p906) sir1Δ::NatNT2 (p994)* |
